# Inducing Specific Chromosome Mis-Segregation in Human Cells

**DOI:** 10.1101/2022.04.19.486691

**Authors:** Laura Tovini, Sarah C. Johnson, Alexander M. Andersen, Diana Carolina Johanna Spierings, René Wardenaar, Floris Foijer, Sarah E. McClelland

## Abstract

Cancer cells display persistent underlying chromosomal instability that results in chromosome mis-segregation, the formation of micronuclei, and abnormal numbers of chromosomes (aneuploidy). These features are common to nearly all human cancers, with individual tumour types intriguingly exhibiting characteristic subsets of whole, and sub-chromosomal aneuploidies. To date, few methods to induce specific aneuploidies at will exist, hampering the investigation of functional consequences of recurrent aneuploidies. Moreover, although some human cell lines with specific aneuploidies exist, the *acute* cellular responses to specific chromosomal instability events remain unknown. We therefore investigated the possibility of sabotaging the mitotic segregation of specific chromosomes using nuclease-dead CRISPR-Cas9 (dCas9) as a cargo carrier to specific genomic loci. We recruited the kinetochore-nucleating domain of centromere protein CENP-T to assemble ectopic kinetochores either near the centromere of chromosome 9, or the telomere of chromosome 1. Ectopic kinetochore assembly led to increased chromosome instability and aneuploidy of the target chromosomes, providing the potential to create ‘designer karyotypes’ and study their immediate downstream cellular responses in a wide range of cell types. Overall, our findings provide new insights into ectopic kinetochore biology, and also represent an important step towards investigating the role of specific aneuploidy and chromosome mis-segregation events in diseases associated with aneuploidy.

## Introduction

Aneuploidy in disease occurs in patterns: Human pluripotent stem cells display recurrent aneuploidies^1^ and congenital aneuploidy syndromes affect a small subset of specific chromosomes^2^. Aneuploidy in cancer is also non-random, with individual cancer types exhibiting characteristic aneuploidy landscapes^3^. In some instances, specific cancer aneuploidies are associated with clinical outcomes, for example the association of monosomy 7 with poor risk acute myeloid leukaemia (AML)^4^. In general however, the functional consequences and clinical implications for specific chromosome alterations remain unknown - in large part due to a lack of tractable cell models that allow investigation into the downstream functional consequences of specific aneuploidies. To date, microcell-mediated chromosome transfer approaches were used to catalogue the impact of specific single chromosome gains and more recently, losses, on cells and have provided important insights into cellular responses to stable expression of a specific aneuploidy^5–7^. Such approaches to manipulate karyotypes rely on a period of selection during which acute responses to aneuploidy may be lost, or adapted to. Meanwhile, CRISPR-targeted telomere cleavage has been utilised to induce chromosome-specific bridges and track the fate of the bridged chromosome in daughter cells using single cell microscopy-based isolation methods^8^. Therefore, the initial cellular responses to gain or loss of specific chromosomes, or other specific chromosome alterations have so far remained elusive. We were thus motivated to discover whether it was possible to induce mis-segregation of a single target chromosome, to allow the study of acute downstream cellular consequences of specific chromosomal alterations.

Elegant prior studies demonstrated that ectopic kinetochores can be induced by artificially recruiting kinetochore-nucleating domains of the inner kinetochore proteins CENP-T, or CENP-C using LacI fusions in cell lines engineered to harbour a LacO array in non-centromeric chromatin^9,10^. CENP-C and CENP-T targeting to LacO arrays initiates the recruitment of downstream kinetochore components, demonstrating the potential to bypass the specialised centromeric CENP-A-containing nucleosomes to form an ectopic kinetochore at non-centromeric locations. Moreover, the resulting pseudo-dicentric chromosomes were subject to faulty chromosome segregation and induced translocations in the LacO-containing chromosome^11^, revealing the potential of this method to induce chromosome mis-segregation. However, several questions remain regarding the functionality of ectopic kinetochores, such as the efficiency of proper kinetochore-microtubule attachment, sister chromatid cohesion, or mitotic checkpoint silencing, and how these functions might vary with genomic position. Moreover, the number of protein moieties required to nucleate a functional ectopic kinetochore could not be answered with the previous systems. In addition, the extremely large and repetitive LacO array employed in these studies carries the potential to form a fragile site^12^ and requires genome editing to create stable cell lines harbouring the LacO array. We therefore designed an approach that could be fine-tuned, allowing the provoked mis-segregation of any specific chromosome, in any given cell line, without the requirement for prior genetic engineering.

Nuclease-dead CRISPR-Cas9 (dCas9) has been extensively used for imaging purposes^13–16^ and also for recruiting functional proteins to the genome, such as transcription factors^17,18^ or chromatin remodellers^19^, as well as centromere protein CENP-B^20^. This prompted us to test whether dCas9 could also nucleate the formation of functional kinetochores at ectopic loci. Such a system would allow the use of endogenous repetitive arrays rather than relying on engineered cell lines harbouring LacO arrays, and the position of the ectopic kinetochore could be varied to target different chromosomes, or chromosomal regions for mis-segregation. Here, we demonstrate that dCas9 can efficiently recruit the kinetochore-nucleating domain of CENP-T (amino acids 1-375, hereafter ‘CENP-T^ΔC^’) to three endogenous repetitive arrays on chromosomes 1, 3 and 9 in both HEK293T and HCT116 human cell lines. Efficient recruitment of outer kinetochore components was achieved to chromosomes 1 and 9, whereupon elevated mis-segregation and aneuploidy of these chromosomes was induced.

## Results

### dCas9 can recruit CENP-T to ectopic loci at comparable levels to endogenous centromeres

We first created CENP-T^ΔC^-dCas9 fusion constructs, including a flexible linker between CENP-T^ΔC^ and dCas9, as previously done when using CENP-T^ΔC^ to nucleate kinetochores^9^. We also fused 3xEGFP proteins to the C-terminus of dCas9 to allow imaging of targeted loci (**Figure 1a**). We expressed the CENP-T^ΔC^-dCas9 fusion protein, using dCas9-EGFP as a control, in HEK293T (human embryonic, transformed) or in HCT116 (human near-diploid colorectal cancer) cells, together with optimised guide RNA scaffolds for increased stability and assembly with dCas9^14^ (**Figure 1c**). We first tested ectopic CENP-T recruitment to the *MUC4* gene, a repetitive locus close to the telomere of chromosome 3 previously used to tether dCas9-EGFP for imaging purposes^14^. Specifically, we targeted a region in the second exon of *MUC4* which contains 100 to 400 repeats of a 48 bp sequence^21^ with a predicted 44-418 binding sites for sgMUC4 (**Figure 1b, Table 1**). Targeting CENP-T^ΔC^-dCas9-EGFP to the *MUC4* locus generated ectopic nuclear foci of CENP-T in the vast majority of transfected (EGFP-positive) HEK293T cells (**Figure 1c,d**). Quantification of signal intensities in metaphase cells revealed that ectopically-recruited CENP-T^ΔC^ levels were on average approximately 60% of CENP-T levels at endogenous centromeres, with individual ectopic CENP-T^ΔC^ foci often overlapping in intensity with lower intensity endogenous centromeres (**Figure 1e**). However, these ectopic CENP-T^ΔC^ foci were not sufficient to recruit the downstream kinetochore component KNL-1 during mitosis (**Figure 1f-h**).

**Figure 1:**
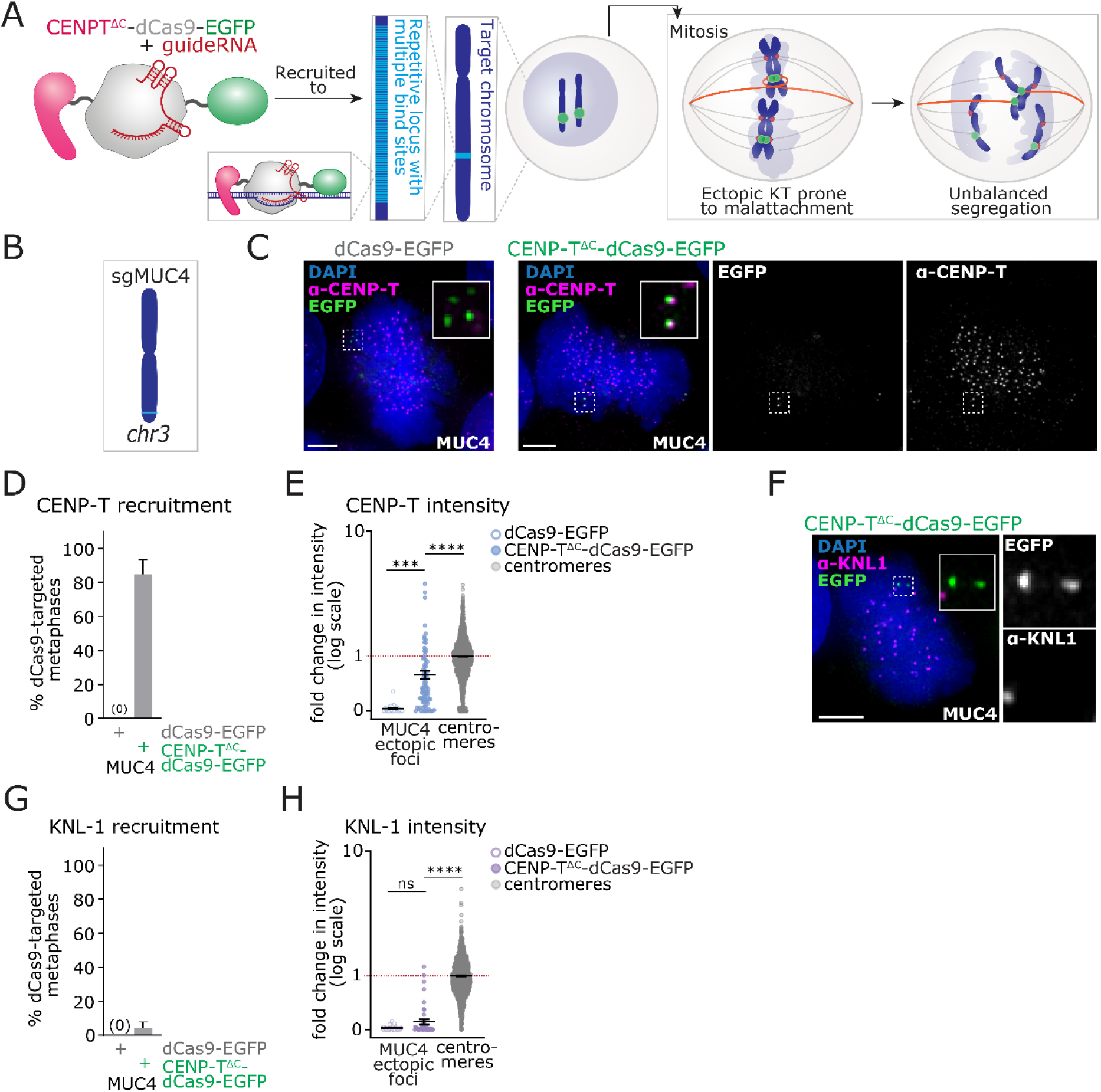
dCas9-mediated recruitment of CENP-T^ΔC^ foci to an ectopic site on Chromosome 3. **A)** Strategy for targeted recruitment of an ectopic kinetochore via dCas9. To specifically recruit CENP-T^ΔC^ to the target chromosome (dark blue), a fusion protein composed of CENP-T^ΔC^ (pink), dCas9 (grey) and EGFP (green) was used with a guide RNA (red) complementary to a sequence (light blue) within a highly repetitive locus. In mitosis, kinetochore-microtubule attachment at endogenous centromeres (red) and ectopic sites (green) renders the target chromosome prone to mis-segregation in anaphase. **B)** Cartoon of chromosome 3 showing the approximate position of *MUC4* target site (light blue). **C-G)** Targeting of dCas9-EGFP or CENP-T^ΔC^-dCas9-EGFP to *MUC4* in HEK293T cells. **C)** Immunofluorescence image of dCas9-EGFP or CENP-T^ΔC^-dCas9-EGFP targeted cells stained with antibodies against CENP-T. **D)** Percentage of metaphase cells showing EGFP and CENP-T signal co-localisation. **E)** Quantification of CENP-T signal intensity at *MUC4* ectopic foci vs. endogenous centromeres, normalised to CENP-T signal intensity at endogenous centromeres (=1, red line). **F)** Immunofluorescence images of dCas9-EGFP or CENP-T^ΔC^-dCas9-EGFP targeted cells stained with antibodies against KNL-1. **G)** Percentage of metaphase cells showing EGFP and KNL-1 signal co-localisation. **H)** Quantification of KNL-1 signal intensity at *MUC4* foci, normalised to KNL-1 signal intensity at centromeres (=1, red line). Data in D and G) Bars = Mean + standard deviation (SD), 3 experiments, each with ≥ 30 metaphases analysed per condition. Data in E and H) Bars = Mean ± SD, 3 experiments, each with ≥ 10 metaphases analysed per condition. Each point = 1 foci or centromere. ns = p>0.05, *** = p<0.01, **** = p<0.001 (Kruskal Wallis test with Dunn’s multiple comparison correction). C and F) Scale bars = 5μm on large images, 1μm on zooms.

**Table 1.**
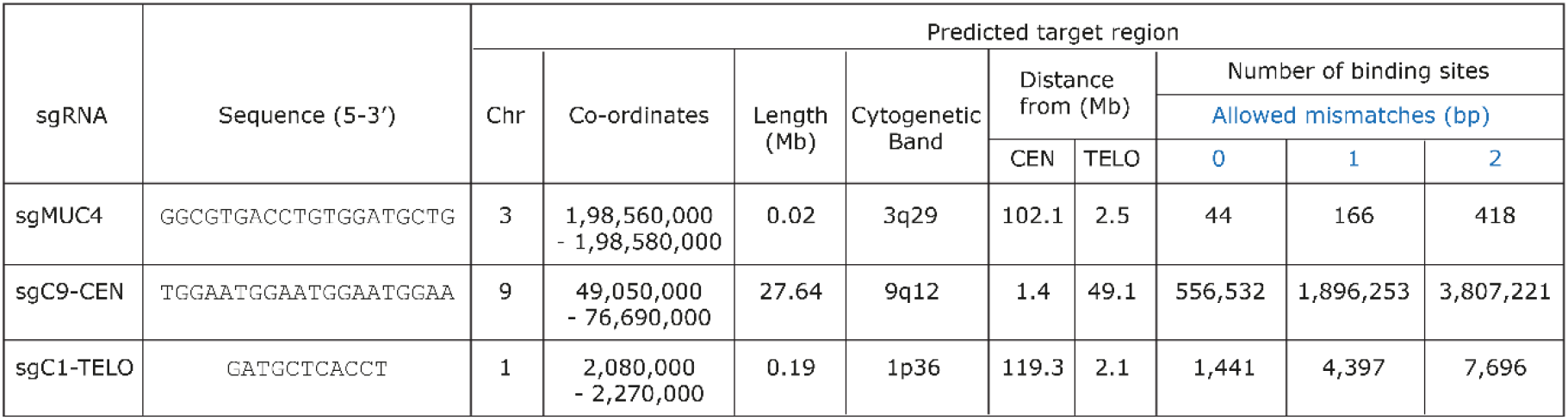
Details of guide RNAs used. For each guide, binding sites within the genome were predicted allowing a range of allowed mismatches between site and guide. Co-ordinates are in CHM13_T2T v1.1 reference genome. CEN = centromere, TELO = closest telomere.

### Increasing the dCas9 binding site size initiates assembly of downstream kinetochore component KNL-1

To test whether recruiting CENP-T^ΔC^-dCas9 to a larger chromosomal target site would allow the establishment of a functional kinetochore, we directed CENP-T^ΔC^-dCas9 to two larger endogenous repetitive arrays. These are located at the pericentromere of human chromosome 9 (‘Chr9-CEN’^13^) or proximal to telomere of the p-arm of human chromosome 1 (‘Chr1-TELO’^20^), and carry a predicted 556,532 – 3.8 million and 1,441 - 7,996 guide RNA binding sites respectively (**Figure 2a; Table 1)**. Fluorescence *In-Situ* hybridisation validated dCas9-EGFP targeting to these loci on metaphase spread chromosomes (**Figure 2b**). Guiding CENP-T^ΔC^-dCas9 to either of these sites resulted in the formation of ectopic CENP-T^ΔC^ foci of significantly higher intensity compared to the *MUC4*-recruited foci in both interphase and mitosis, in both HEK293T and HCT116 cells (**Figure 2c-e; Figure S1a,b**) (the presence of three signals for Chr9-CEN is due to the near-triploid karyotype of HEK293T cells^22^). Strikingly, KNL-1 was now efficiently recruited to both Chr9-CEN and Chr1-TELO in both cell lines during prometaphase when CENP-T-mediated KNL-1 loading usually occurs^23^ (**Figure 2f-h; Figure S1c,d**), resulting in ectopic KNL-1 foci of 3-fold (Chr1-TELO), or 10-fold (Chr9-CEN) higher intensity compared to endogenous centromeres (**Figure 2h**). Ndc80 was also efficiently recruited to Chr1-TELO and Chr9-CEN sites (**Figure 2i,j; Figure S1e,f**).

**Figure 2:**
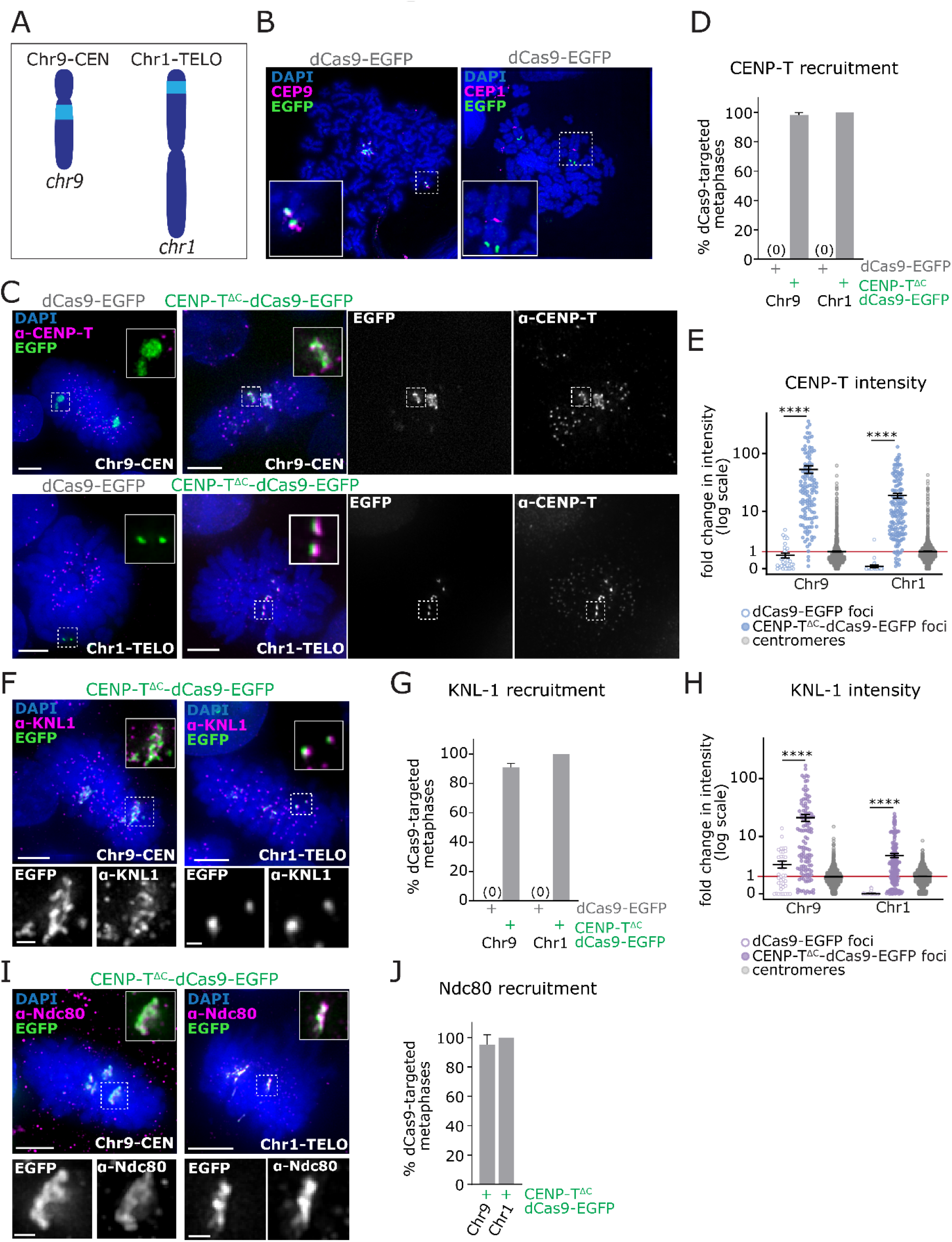
CENP-T^ΔC^-dCas9 targeting recruits KNL-1 and Ndc80 to large repetitive chromosomal loci. **A)** Cartoon showing the approximate position of the Chr9-CEN and Chr1-TELO target sites (light blue). **B-I)** Targeting of dCas9-EGFP or CENP-T^ΔC^-dCas9-EGFP to Chr9-CEN and Chr1-TELO in HEK293T cells. **B)** FISH images on chromosome spreads from cells with dCas9-EGFP targeting. **C)** Immunofluorescence images of dCas9-EGFP or CENP-T^ΔC^-dCas9-EGFP targeted cells stained with antibodies against CENP-T. **D)** Percentage of metaphase cells showing EGFP and CENP-T signal co-localisation. **E)** Quantification of CENP-T signal intensity at Chr9-CEN and Chr1-TELO ectopic foci vs. endogenous centromeres, normalised to CENP-T signal intensity at endogenous centromeres (=1, red line). **F)** Immunofluorescence images of CENP-T^ΔC^-dCas9-EGFP targeted cells stained with antibodies against KNL-1. **G)** Percentage of metaphase cells showing EGFP and KNL-1 signal co-localisation. **H)** Quantification of KNL-1 signal intensity at Chr9-CEN and Chr1-TELO foci vs. endogenous centromeres, normalised to KNL-1 signal intensity at centromeres (=1, red line). **I)** Immunofluorescence images of dCas9-EGFP or CENP-T^ΔC^-dCas9-EGFP targeted cells stained with antibodies against Ndc80. **J)** Percentage of metaphase cells showing EGFP and Ndc80 signal co-localisation. Data in D,G and J) Bars = Mean + SD, 3 experiments, each with ≥50 metaphases per condition. Data in E and H) Bars = Mean ± SD, 3 experiments, each with ≥10 metaphases per condition. ns = p>0.05, **** = p<0.001 (Kruskal Wallis test with Dunn’s multiple comparison correction). B,C,F and I) Scale bars = 5μm on large images, 1μm on zooms.

### Ectopic kinetochores attach to, and stabilise microtubules

Similar to ectopic kinetochores generated with the LacO system^9^, mitotic but not interphase CENP-T^ΔC^-dCas9 foci recurrently showed a bar-like shape, whereas dCas9-EGFP control foci remained circular during mitosis (**Figure 3A**). This suggested interaction with mitotic spindle microtubules, and accordingly, these bar-like CENP-T^ΔC^-dCas9 foci reverted to circular morphology upon nocodazole-induced microtubule depolymerisation (**Figure 3B**). We therefore examined CENP-T^ΔC^-dCas9 foci for the presence of stably attached spindle microtubules as an indication of the functionality of the ectopic kinetochore. Cells were cold treated for 10 minutes to depolymerise non-kinetochore attached microtubules^24^. The majority of mitotic cells exhibiting ectopic kinetochores at either chromosome 1, or chromosome 9 displayed obvious kinetochore fibres terminating at the ectopic CENP-T^ΔC^/KNL-1 site which were often abnormally large at Chr9-CEN (**Figure 3C,D)**. Such fibres or bundles were rarely observed in EGFP-negative (untransfected) cells or dCas9-EGFP targeted cells. Taken together these data suggest that dCas9-tethered CENP-T^ΔC^ is able to recruit functionally active downstream kinetochore components and provide attachment to mitotic spindle microtubules.

**Figure 3:**
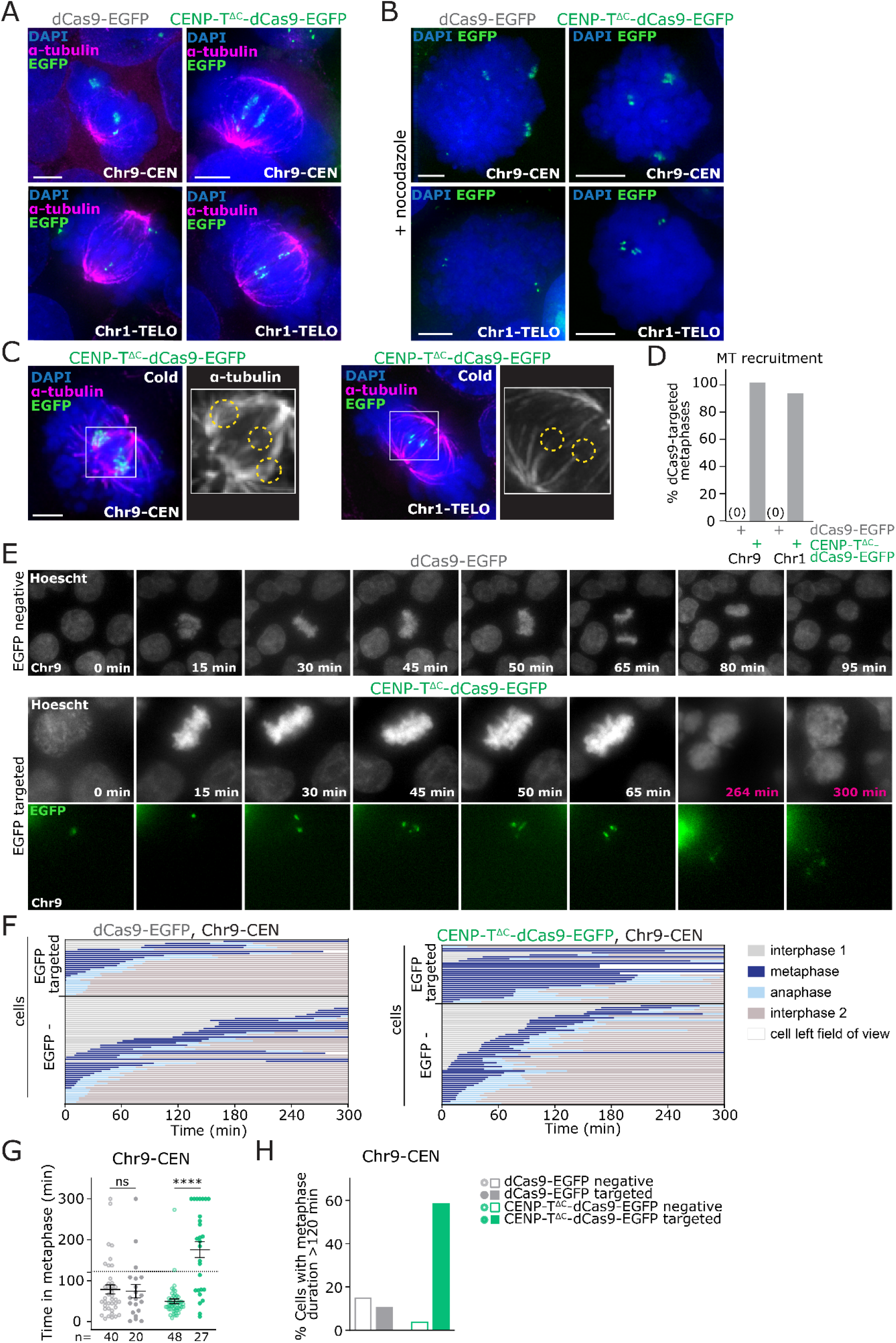
CENP-T^ΔC^-dCas9-nucleated ectopic kinetochores recruit microtubules but induce mitotic delay. **A-H)** Targeting of dCas9-EGFP or CENP-T^ΔC^-dCas9-EGFP to Chr9-CEN and Chr1-TELO in HEK293T cells. **A)** Immunofluorescence images of dCas9-EGFP or CENP-T^ΔC^-dCas9-EGFP targeted metaphase cells stained with antibodies against alpha-tubulin. **B)** Immunofluorescence images of mitotic cells with dCas9-EGFP targeting after nocodazole treatment. **C)** Immunofluorescence images of metaphase cells with CENP-T^ΔC^-dCas9-EGFP targeting after cold treatment, stained with antibodies against alpha-tubulin. **D)** Percentage of metaphase cells showing microtubule (MT) recruitment. One experiment, ≥50 metaphases per condition. **E)** Frames from live cell imaging of cells with dCas9 targeted to Chr9-CEN, or an EGFP negative cell. **F)** Cell cycle stages of cells with CENP-T^ΔC^-dCas9-EGFP or dCas9-EGFP transfected cells, by live-cell imaging (each line = 1 cell, ≥ 27 cells per group), showing EGFP negative and EGFP targeted cells separately. **G)** Time spent in metaphase for CENP-T^ΔC^-dCas9-EGFP or dCas9-EGFP transfected cells, showing EGFP negative and EGFP targeted cells separately. Dotted line = 120 min = cut-off for the longest time that the majority of control cells spent in metaphase. **H)** Percentage of cells that spent ≥120 min in metaphase. Data in F-H) are from one experiment, Bars = mean (± Standard error of the mean, SEM). ns = p>0.05, **** = p<0.001 (One-way ANOVA with Šidák’s multiple comparison correction).

### CENP-T^ΔC^-nucleated kinetochores trigger the mitotic spindle assembly checkpoint

The expected outcome of an ectopic kinetochore is the creation of a ‘pseudo-dicentric’ chromosome which has an increased chance of becoming incorrectly attached to microtubules and is therefore more prone to faulty mis-segregation^11^ (see schematic in **Figure 1a**). To test if this would occur with dCas9-nucleated ectopic kinetochores, we performed live cell imaging of HEK293T cells with dCas9-CENP-T^ΔC^ targeted to Chr9-CEN (**Figure 3e; Movie S1**). Surprisingly, most mitotic cells exhibited a prolonged mitotic delay; the average length of metaphase in cells with dCas9-CENP-T^ΔC^ ectopic kinetochores was 175 minutes compared with 74 minutes in dCas9-EGFP targeted cells and 50 minutes in EGFP-negative cells (**Figure 3f-h**). Since cells with ectopic kinetochores were often already in metaphase at the start of the movie this was also an underestimation of the metaphase delay. Fixed cell imaging confirmed the metaphase delay for both Chr9-CEN and Chr1-TELO (**Figure 4a,b; Figure S2a,b**).

**Figure 4:**
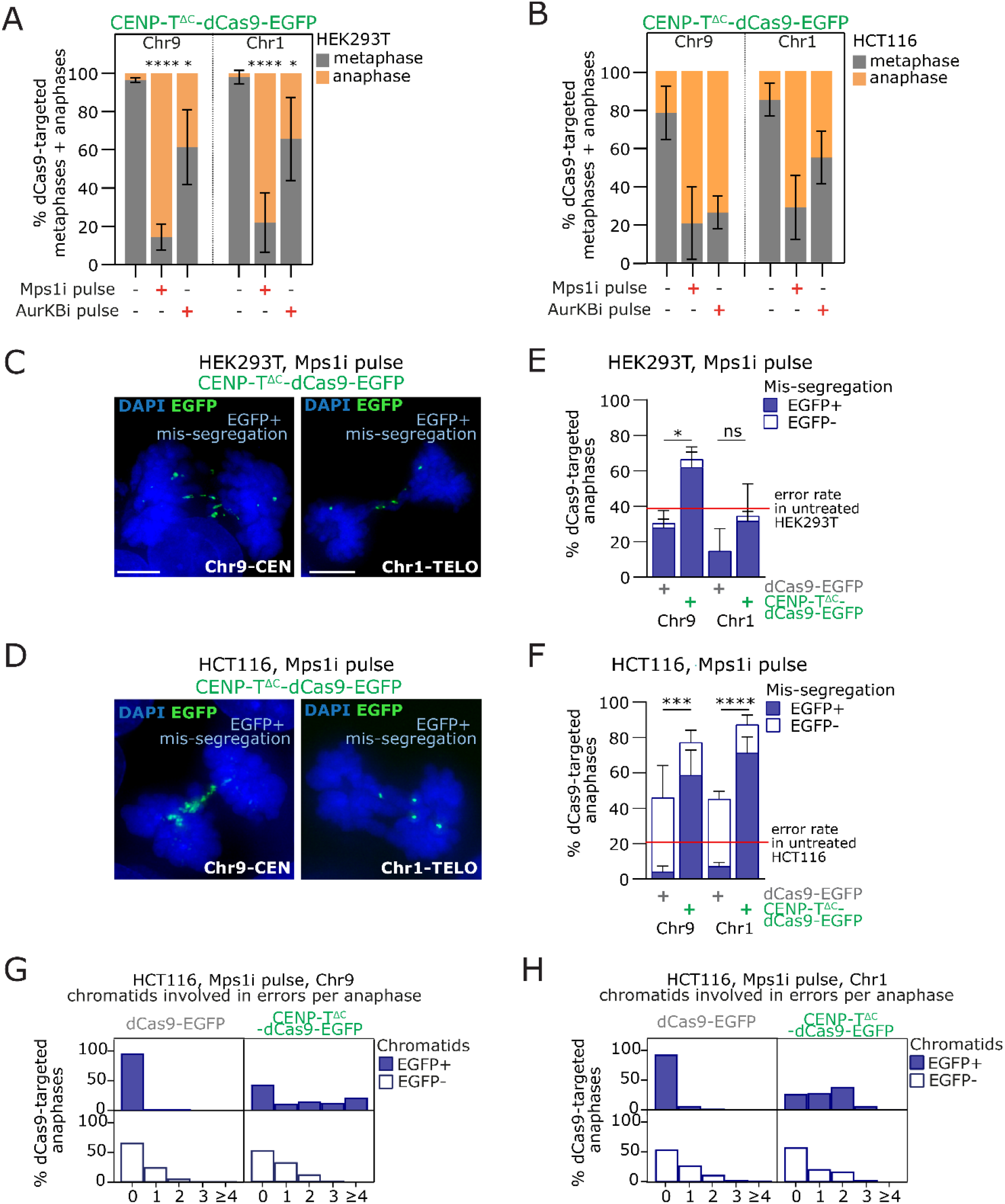
Induction of specific chromosome mis-segregation by ectopic kinetochore strategy. **A,B)** Quantification of mitotic stage from fixed CENP-T^ΔC^-dCas9 targeted HEK293T (**A**) or HCT116 (**B**) cells following inhibition of Mps1 (Mps1i) or Aurora Kinase B (AurKBi) (Bars = mean ± SD, 3 experiments for HEK293T, 2 for HCT116, each with ≥50 mitotic cells per condition). **C,D)** Example images of chromosome mis-segregation events involving the target locus (EGFP signal) in anaphase HEK293T (**C**) or HCT116 (**D**) cells with CENP-T^ΔC^-dCas9-EGFP targeting after Mps1i pulse. **E,F)** Mis-segregation rate in dCas9-targeted HEK293T (**E**) or HCT116 (**F**) cells after Mps1i pulse, either involving EGFP+ chromosomes, or only EGFP-chromosomes. Red line = mean error rate in untreated cells. **G,H)** Quantification of number of EGFP+ (**G**) or EGFP- (**H**) chromatids involved in segregation errors from HCT116 anaphases with dCas9 targeting. Statistics in A,B,E and F) ns = p>0.05, * = p<0.05, *** = p<0.001, **** = p<0.0001 (One-way ANOVA with Šidák’s multiple comparison correction, in E/F) comparing the EGFP+ mis-segregation rate).

### CENP-T^ΔC^-nucleated ectopic kinetochores induce a mitotic arrest via Aurora B activity

The observed metaphase arrest suggested an activated mitotic checkpoint. Accordingly, treating CENP-T^ΔC^-dCas9 or dCas9-EGFP-targeted cells with Mps1 kinase inhibitor NMS-P715^25^ to abrogate the mitotic checkpoint^26^ allowed the majority of metaphase-arrested cells to progress into anaphase (**Figure 4a,b; Figure S2a,b**). We reasoned that Aurora B mediated detachment of improperly attached ectopic kinetochores could underlie the activation of the mitotic checkpoint. To test this, we inhibited Aurora B in transfected cells using ZM447439^27^. Aurora B inhibition reduced the metaphase arrest seen upon targeting CENP-T^ΔC^-dCas9 to either Chr9-CEN or Chr1-TELO, matching the efficiency of Mps1 inhibition in HCT116 cells (**Figure 4a,b; Figure S2a-c**). This suggests that a large proportion of metaphase arrest is caused by improper ectopic kinetochore attachments that activate the mitotic checkpoint in a manner dependent on Aurora B-mediated error correction. In some cells however, inhibition of Aurora B could not overcome the metaphase arrest (particularly in HEK293T cells).

### CENP-T^ΔC^-nucleated ectopic kinetochores induce specific mis-segregation of targeted chromosomes

We reasoned that abrogating the mitotic arrest by inhibiting the Mps1 kinase^26^ would allow us to score the impact of ectopic kinetochore assembly on chromosome segregation fidelity. We therefore scored CENP-T^ΔC^-dCas9-targeted chromosome segregation error rates from anaphase cells that were visible after a short Mps1i pulse, exploiting the EGFP fluorescence present on ectopic kinetochores to detect the target chromosome in the act of mis-segregation. We classified as ectopic kinetochore mis-segregation any improper segregation event where EGFP signals were observed lagging in the centre of the segregating anaphase DNA masses (**Figure 4c,d**). Cells with ectopic KT mis-segregation occasionally also mis-segregated non-EGFP chromosomes. In HCT116 cells we were able to quantify the numbers of GFP-negative or positive mis-segregating chromatids per cell (**Figure 4g,h**). In HEK293T cells, which could not be analysed in this way due to less distinct chromosome mis-segregation events, we simplified the analysis to cells with EGFP+ mis-segregation, or non EGFP+ chromosome mis-segregation (**Figure 4e,f**). Cells with ectopic kinetochores displayed higher EGFP+ chromosome mis-segregation rates compared to dCas9-EGFP alone controls (**Figure 4e,f**). This was true for both chromosome 9 and 1, and in both HEK293T and HCT116 cells (**Figure 4c-f**).

### Induction of aneuploidies on target chromosomes

To directly test the consequences of targeted mis-segregation on aneuploidy of daughter cells we sequenced individual G1 cells following transfection and a pulse of Mps1i. Cells were allowed to complete mitosis and separate into two daughter cells before undergoing single cell low pass whole genome sequencing. HEK293T cells contain a rearranged, but stable karyotype with only minor translocations involving chromosome 1 or 9^22^. We therefore computed copy number gains and losses compared to a pseudo bulk HEK293T reference obtained from our control (Mps1i, no guide no dCas9) population of sequenced cells (**Figure 5a; Figure S3a**, see methods). ClonalMasker (see methods) was used to remove clonal and subclonal aneuploidies (present in identical positions across more than one cell) present in the Mps1i only control from each additional condition, and report back only unique aneuploidies >20Mb in length (see pileups in **Figure 5b,d; Figure S3b**). We then scored partial and whole chromosome aneuploidy events > 20 Mb for each chromosome, across all conditions (**Figure 5c,e,f; Figure S3c**). Determining new aneuploidy events occurring on a heterogeneous background is challenging, and in addition only approximately 40% cells displayed CENP-T^ΔC^-dCas9 targeting within this experiment. Nonetheless, with CENP-T^ΔC^ targeting to chromosome 1 and 9 we were able to detect an enrichment in aneuploidies affecting the target chromosomes (**Figure 5b-f**). Interestingly the dCas9-EGFP control targeting to chromosome 1 also induced aneuploidy of chromosome 1 although we interpret this result with caution since chromosome 1 exhibited a high frequency of aneuploidies across most conditions likely due to additional sub-clonal aneuploidies present in the parental line (**Figure 5e**). Closer examination of induced CNAs (**Figure 5b,d**) revealed a specific enrichment of breakpoints proximal to the target sites. For CENP-T^ΔC^ targeting to Chr9-CEN (**Figure 5b**), induced CNAs were all found with breakpoints in the q-arm proximal to the peri-centromeric target site. Meanwhile, ectopic kinetochore targeting to Chr1-TELO (**Figure 5d**) led to an enrichment of breakpoints between the target telomere and endogenous centromere but at a larger distance from the actual target site, suggesting physical breakage of the chromosome between two opposing attachment sites as expected for a dicentric chromosome. Taken together these data show we can use dCas9 to induce specific chromosome mis-segregation and aneuploidy, and that the strategic use of different chromosomal locations as ectopic kinetochore sites can be used to induce specific types of chromosome segregation error.

**Figure 5:**
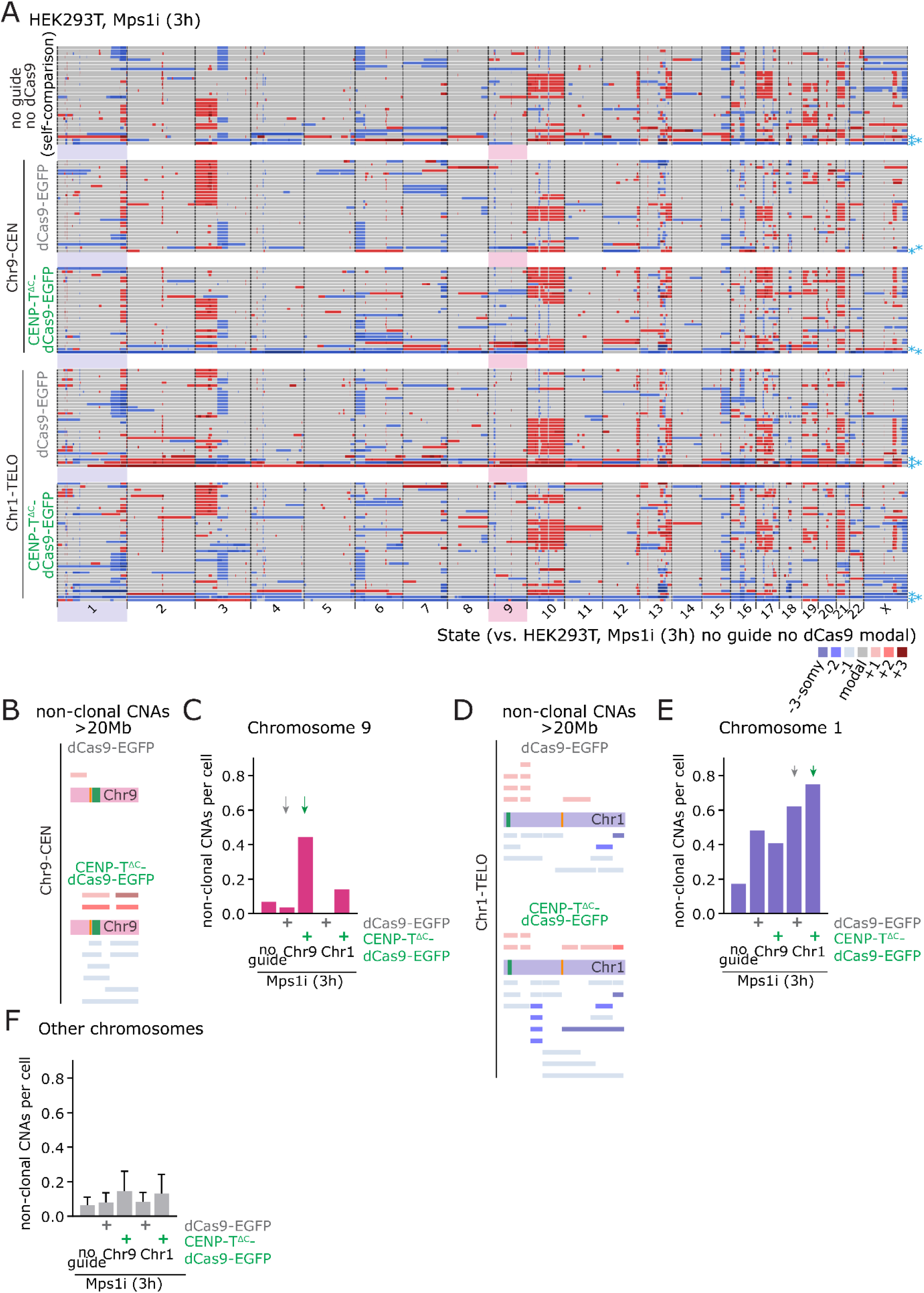
Single cell sequencing reveals chromosome-specific aneuploidies induced by CENP-T^ΔC^-dCas9-nucleated kinetochores. **A)** Copy number calls from single cell sequencing data. Colours indicate copy number relative to the modal karyotype of control HEK293T cells treated with Mps1i (no guide, no dCas9), in the conditions indicated. * = cells excluded from further analysis due to the presence of widespread aneuploidy relative to the median reference genome. Shaded boxes = target chromosomes. 28 - 39 cells sequenced per condition **B-F)** Assessment of non-clonal CNAs > 20Mb. Any clonal CNAs, and subclonal CNAs observed in the control (HEK293T, Mps1i (3h), no guide no dCas9) were omitted. **B and D**) CNA pileups showing location of CNAs in target chromosomes, for chromosome 9 (B) and chromosome 1 (D). Orange lines indicate endogenous centromeres and green lines indicate CENP-T^ΔC^ target sites. **C, E and F**) Quantification of non-clonal CNAs >20Mb affecting chromosomes 1 (C) or 9 (E), or other chromosomes (F). Frequency of aneuploidy events >20 Mb per chromosome per cell is indicated. Bars = mean (+ SD). Arrows highlight conditions shown in pileups.

## Discussion

In this study we induced the specific mis-segregation of human chromosomes using dCas9 to nucleate ectopic kinetochores at endogenous repeat arrays. Large endogenous repetitive arrays in chromosomes 1 and 9 allowed the efficient recruitment of ectopic kinetochores that bound microtubules and caused elevated mis-segregation and aneuploidy rates of those chromosomes. In an accompanying manuscript (Truong et al., BIORXIV 2022), tethering of a plant kinesin using TetR repeats, or dCas9 similarly induced elevated mis-segregation of targeted chromosomes, demonstrating the flexibility with which dCas9 fusion proteins can be used to interfere with chromosome segregation. Together our studies provide an important step towards custom manipulation of mitosis to induce specific aneuploidies.

### What length and position of target array is required for ectopic kinetochore formation?

The smallest array (the *MUC4* locus on chromosome 3), despite recruiting CENP-T^ΔC^ to comparable levels as lower intensity endogenous centromeres, was unable to assemble an ectopic kinetochore. Instead, arrays providing upwards of 1441 guide RNA binding sites were able to form functional kinetochores, providing the boundaries for the length of target arrays required to assemble CENP-T-nucleated ectopic kinetochores, though we did not determine the precise threshold in this study. In terms of positioning, we were able to provoke mis-segregation using both centromere-, and telomere-proximal sites, however it is possible that the exact underlying causes of mis-segregation may vary between these two positions. For example, CEN9-targeting might interfere with the endogenous chromosome 9 centromere function, potentially mimicking merotelic attachment, while TELO1-targeting might be more likely to create a canonical pseudo-dicentric chromosome. Customising the location of ectopic kinetochore formation would therefore theoretically allow the modelling of specific cancer-associated events, such as arm-level aneuploidy generation, improper kinetochore-microtubule attachments, and dicentric chromosomes.

### CENP-T nucleated kinetochores activate the mitotic checkpoint

Despite building a functional microtubule attachment site, and apparently aligning at the metaphase plate, CENP-T-nucleated kinetochores could not readily satisfy the mitotic checkpoint. Aurora B inhibition completely (HCT116), or partly (HEK293T) overcame the mitotic arrest, potentially due to suppression of improper kinetochore-microtubule detachment. Aurora B’s role in establishment of the mitotic checkpoint *per se*^28^ means this result should be interpreted with caution, although cells were arrested in metaphase with a functional checkpoint established at the time of treatment. In addition, it is possible that cells that failed to undergo anaphase following inhibition of Aurora B were unable to correctly strip mitotic checkpoint proteins from ectopic kinetochores. Alternatively, this could be due to unattached ectopic kinetochores, although we did not notice the presence of obviously unattached ectopic kinetochores herein. It will be interesting to characterise these pathways further and in different chromosomal and cellular contexts.

### Future development of dCas9-based strategies to induce specific chromosome mis-segregation

Herein, we used transient transfections in HEK293T and HCT116 cells to allow rapid optimisation of the system, and to avoid the insertion of exogenous genomic material, therefore providing proof-of-principle of how specific chromosome segregation defects can be induced *ad hoc* in cell lines without any prior genetic editing required. However, we anticipate that generation of stable cell lines with the capacity to create specifically targeted ectopic kinetochores at any locus, determined by the transfection with specific guide RNAs will provide a powerful model system to explore the cellular and genomic consequences of specific chromosome mis-segregation.

## Availability of data and materials

All raw sequencing data will be accessible via the European nucleotide database upon publication

## Author contributions

LT and SCJ designed and performed all cell biological experiments and data analysis supervised by SEM. RW and DCJS performed and analysed single cell sequencing data supervised by FF. AA performed bioinformatics supervised by SEM. SEM conceived the study, designed experiments, analysed data and wrote the manuscript with input from all authors.

## Acknowledgements

We would like to thank Susanne Lens, My Anh Truong, Paula Cané-Gasull and Sippe de Vries for helpful discussions and sharing of unpublished data and reagents. We would also like to thank Iain Cheeseman for constructs (via Addgene) and constructive discussions, Steve Royle for discussions and advice and Victoria Sanz-Moreno for HEK293T cells. We would also like to dedicate this work to the memory of Norah Reed, founder of the Barry Reed Charity, whose generosity and genuine interest in the scientific findings and career progression of LT was an inspiration to us.

## Methods

### Cell culture and drug treatments

Cells were grown in DMEM (41966; Thermo Fisher Scientific) supplemented with 10% (v/v) Gibco™ Foetal Bovine Serum (FBS) (10500064; Thermo Fisher Scientific) and 1% (v/v) Penicillin-Streptomycin (#P4333; Sigma Aldrich) at 37 °C and 5% CO_2_. HEK293T cells were a gift from Prof. Victoria Sanz-Moreno, and HCT116 cells were a gift from Prof. Charles Swanton. Routine STR (Public Health England) and mycoplasma checks (#LT07-118; Lonza) were conducted to ensure cell line identification and mycoplasma-free status. For drug pulses (**Figure 4; Figure S2**), cells were treated with 10μM Aurora B inhibitor (ZM447439, Cayman chemical) or 1μM Mps1 inhibitor (NMSP715; Sigma Aldrich) for 20min for HCT116 and 30min for HEK293T as was determined to be sufficient to allow metaphase-arrested cells to reach anaphase. For single cell sequencing (**Figure 5; Figure S3**), a longer Mps1i treatment of 3h was used. For microtubule depolymerisation experiments (**Figure 3b**), cells were treated with 100ng/ml Nocodazole for 4h.

### Plasmids

For the generation of the CENP-T^ΔC^-dCas9-EGFP plasmid, the coding sequence for the CENP-T^ΔC^ fragment^6^ was amplified from an existing plasmid (#45109; Addgene) for insertion into a dCas9-EGFP containing backbone (pHAGE-TO-dCas9-3XEGFP (#64107; Addgene)). In the amplification PCR for CENP-T^ΔC^, coding sequences for flexible linkers of 2xGlycine_4_Serine were added on the primers such that they would flank CENP-T^ΔC^ in the fusion protein. For sgMUC4 and sgC9-CEN, a custom plasmid backbone was constructed by inserting an optimised guide RNA scaffold^14^ under a U6 promoter into the mCherry-C1 plasmid (#54563; Addgene). Proceeding the guide RNA scaffold a BsmBi-excisable sequence was included to allow for targeting sequence introduction as per the “Lentiviral CRISPR/Cas9 and single guide RNA” protocol from the Zhang lab (Joung et al., 2017). sgChr1-TELO was a kind gift from Prof. Susanne Lens^20^. Targeting sequences for each guide RNA are detailed in **Table 1**.

### Plasmid transfection

Plasmid expression was achieved by transient transfection of cells 24h after plating with a dCas9:guide RNA plasmid ratio of 3:1. The DNA solution and the transfection reagent (Lipofectamine 2000; Thermo Fisher Scientific) were separately prepared in Opti-MEM™ media (#31985062; Gibco) and mixed following manufacturer’s instructions. Transfection was performed by adding the transfection mix dropwise on the cell culture plates and cells harvested for further analysis 16-24h later.

### Immunofluorescence

Cells grown on glass slides or coverslips were fixed with PTEMF (0.2% Triton X-100, 0.02 M PIPES [pH 6.8], 0.01 M EGTA, 1 mM MgCl2, and 4% formaldehyde). After blocking with 3% BSA PBS for 1h at 21°C, cells were incubated with primary antibodies in blocking buffer for 1h at 21°C: 1:1000 α-tubulin (118M4779V; Sigma Aldrich), 1:250 CREST (15-234-0001; Antibodies Incorporated), 1:500 CENP-T (ab220280; Abcam), 1:500 KNL-1 (ab222055; Abcam), 1:500 Mad2 (ab97777; Abcam), 1:500 Ndc80 (ma1-213308; Invitrogen). For Ndc80 staining, blocking was conducted instead in 5% Milk PBS. Secondary antibodies used were goat anti-human AF647 (109-606-088-JIR; Stratech), goat anti-rabbit (A11012; Invitrogen), goat anti-mouse (A11005; Invitrogen) at 1:250 dilution. DNA was stained with DAPI (Roche), and coverslips mounted in Vectashield (Vector H-1000; Vector Laboratories). Following each antibody incubation, cells were washed in PBS three times, each for 3 min.

### Metaphase Spreads

To enrich for mitotic cells, cells were treated with 0.1 μg/ml Colcemid (#15212012; ThermoFisher Scientific) for 2h before conducting mitotic shake-off. Collected cells were pelleted and re-suspended in 75 mM KCl hypotonic solution for 10-15 min depending on cell type, on ice. Cells were pelleted and re-suspended in freshly prepared 3:1 methanol-glacial acetic acid a total of 3 times, before dropping onto slides.

### Fluorescence *In Situ* Hybridization

Slides with freshly made metaphase spreads were put through an ethanol dehydration series and air dried. Cells and specific centromere probe (LPE 001R or LPE 009R; Cytocell) were denatured for 2 min at 75 °C then incubated overnight at 37 °C. The following day, slides were washed with 0.25X SSC at 72 °C followed by a brief wash in 2X SSC, 0.05% Tween. DNA was stained for 6 min with 0.5 μg/ml DAPI (Roche) and coverslips mounted in Vectashield (Vector H-1000, Vector Laboratories).

### Microscopy

Images were acquired using an Olympus DeltaVision RT microscope (Applied Precision, LLC) equipped with a Coolsnap HQ camera and CO2 controlled chamber. Three-dimensional image stacks were acquired at 0.2 μm intervals, using Olympus 100X (1.4 numerical aperture) UPlanSApo oil immersion objectives. Deconvolution of image stacks was performed with SoftWorxExplorer (Applied Precision, LLC). For live-cell imaging, HEK293T cells were seeded in four well imaging dishes (Greiner Bio-one) and imaging begun 16h following transfection. Prior to imaging, cells were stained with 0.05 μg/ml Hoechst DNA staining solution (Thermo Fisher Scientific) for 10 min, then washed three times and imaged in freshly added media. Cells were imaged using an Olympus 40X 1.3 numerical aperture UPlanSApo oil immersion objective, with image stacks at 20 μm intervals (10 images) taken every 3 min for 5h. Analysis was performed using Softworx Explorer.

### Quantification of CENP-T and KNL-1

The intensity of the CENP-T and KNL-1 fluorescence signal was measured using Imaris software (Bitplane; version9.1). For each cell, the targeted EGFP signal and all endogenous centromeres were marked using automated detection, using spot-based detection based on anti-centromere serum signal for each centromere, and surface-based detection for EGFP. Intensity measurements were taken for the channel of interest (either CENP-T or KNL-1) at these marked regions (targeted EGFP and endogenous centromeres). After subtracting average background intensity, intensity values were then normalised by dividing the average intensity at the EGFP focus by the average centromere intensity (using values from the same cell).

### Estimation of guide RNA binding sites abundancy

Guide RNA binding site predictions were run using the command line version of Cas-OFFinder algorithm^29^ against the T2T-CHM13 (version 1.1) reference genome^30^. For generation of plots, chromosome reference sequences were binned into 10Kbp lengths, and any adjacent bins with one or more binding sites predicted were then merged to form a single binding cluster. The number of predicted binding sites was then calculated per cluster, and the details of the cluster with the most binding sites reported (**Table.1**). Genome binning and cluster analysis was conducted in R (version 4.1.1, R Foundation for Statistical Computing) using RStudio interface (RStudio).

### Single cell sequencing and analysis

Single G1 nuclei, as assessed by PI and Hoechst staining, were isolated by flow cytometric cell sorting into 96-well plates and preamplification-free single-cell whole genome sequencing libraries prepared using a Bravo automated liquid handling platform (Agilent Technologies) as previously described^31,32^. In brief, genomic DNA was fragmented using micrococcal nuclease followed by end-repair, A-tailing and Illumina PE forked adapter ligation. Upon AMPure XP bead clean-up, the adapter-containing DNA fragments were subjected to PCR amplification using multiplexing primers to incorporate library-specific barcodes. Pooled libraries were subsequently shallow sequenced on an Illumina NextSeq 500 sequencer (up to 77 cycles; single end) with ^~^ 1% genomic DNA coverage. The generated data were demultiplexed using library-specific barcodes and changed into fastq files using bcl2fastq (Illumina; version 1.8.4). Reads were afterwards aligned to the human reference genome (GRCh38/hg38) using Bowtie2 (version 2.2.4^33^). Duplicate reads were marked with BamUtil (version 1.0.3^34^).

Copy number calls were made using AneuFinder (version1.14.0^35^). To account for the high level of existing heterogeneity seen in controls, modal copy number calls were calculated by taking the median copy number across the individual cells (HEK293T, Mps1i (3h), no guide no dCas9; **Figure S3A**). Copy number states for the single cells from other conditions were then recalculated relative to this (**Figure 5A**). ClonalMasker was used to filter out CNAs that occurred in more than 1 cell and were therefore considered clonal (or sub-clonal), for the control sample (Mps1i). A minimum overlap of 80% was utilised, and a fraction that secured only CNAs present in 1 cell was called (1/29 = 0.035). Following this cleaning of the control, the control CNAs were used to call CNAs from the test conditions (with dCas9-EGFP/CENP-T^ΔC^-dCas9-EGFP targeted to Chr9-CEN or Chr1-TELO) that were not present in the control. Analysis was carried out in Excel and R, where CNAs > 20 Mb were isolated. Pileup graphs were created by running the unique CNAs through ClonalMasker with a frequency of 1.0, allowing for all CNAs to be plotted. Clonal CNA filtering scripts are available at https://github.com/MBoemo/clonalMasker.

### Preparation of Data and Figures

All graphs were prepared, and statistical testing performed in Prism software (version 9.0, GraphPad). Contrast and brightness of the final images were linearly adjusted in Photoshop (Adobe Photoshop CS6 2018, USA). All figures were assembled in Illustrator (Adobe Illustrator CS6 2018).

---

**Supplementary Figure 1.**
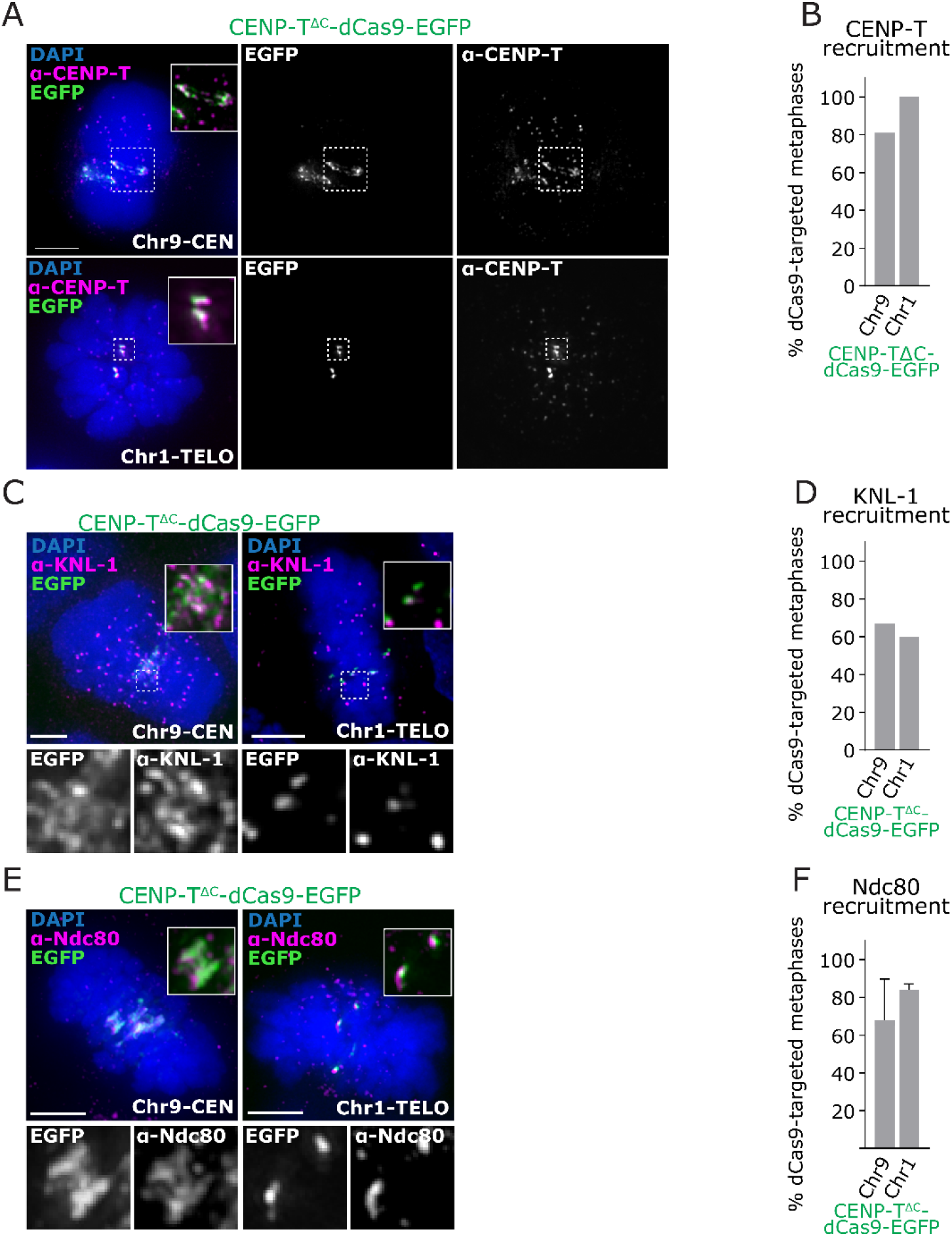
related to Figure 1. **A)** Immunofluorescence images of HCT116 cells with CENP-T^ΔC^-dCas9-EGFP targeted to Chr9-CEN or Chr1-TELO, stained with antibodies against CENP-T. **B)** Percentage of metaphase cells showing EGFP and CENP-T signal co-localisation. ≥20 metaphases analysed per condition, 1 experiment. **C)** Immunofluorescence images of HCT116 cells with CENP-T^ΔC^-dCas9-EGFP targeted to Chr9-CEN or Chr1-TELO, stained with antibodies against KNL-1. **D)** Percentage of metaphase cells showing EGFP and CENP-T signal co-localisation. ≥20 metaphases analysed per condition, 1 experiment. **E)** Immunofluorescence images of HCT116 cells with CENP-T^ΔC^-dCas9-EGFP targeted to Chr9-CEN or Chr1-TELO, stained with antibodies against Ndc80. **F)** Percentage of metaphase cells showing colocalization of EGFP with Ndc80. ≥20 metaphases counted per condition, 2 experiments, error bars = SD.

**Supplementary Figure 2.**
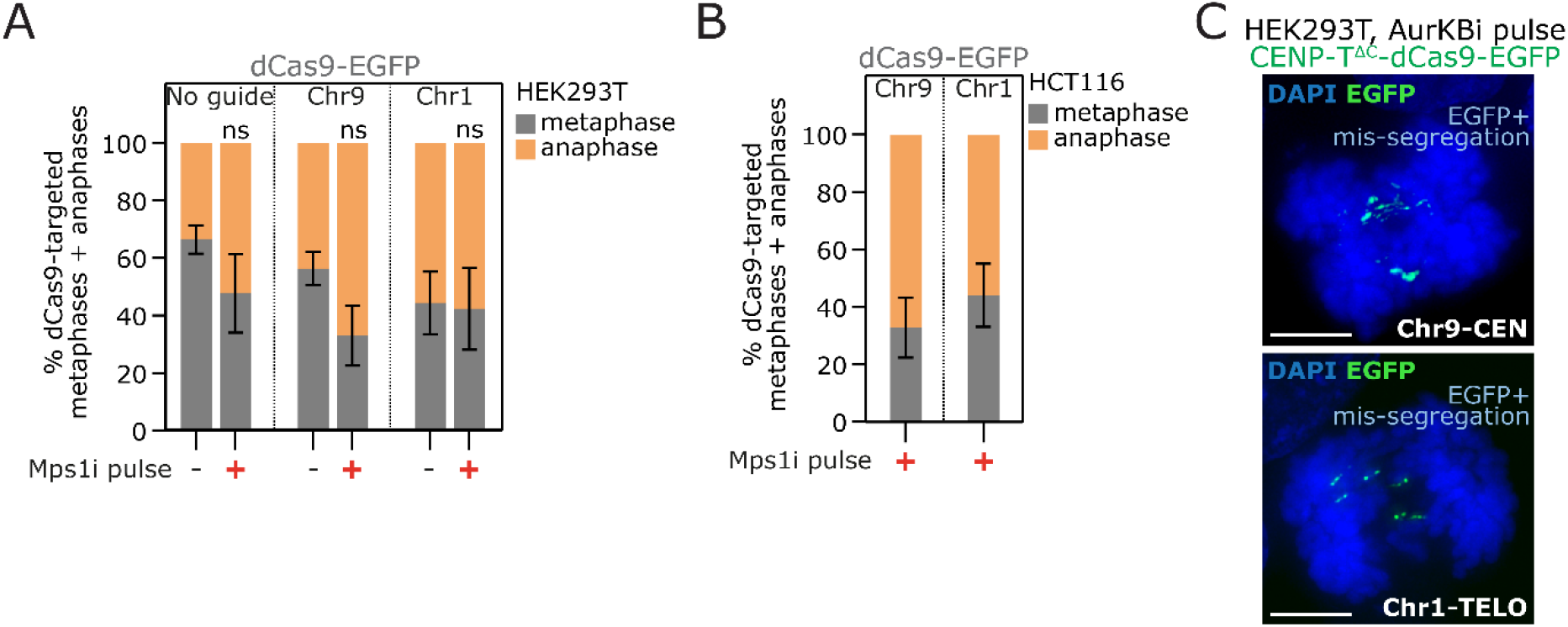
related to Figure 4. **A-B)** Quantification of mitotic stage from fixed dCas9-EGFP targeted HEK293T (**A**) or HCT116 (**B**) cells following short inhibition of Mps1 (Mps1i). Bars = mean ± SD, ≥50 mitotic cells analysed per condition, 3 experiments. **C)** Immunofluorescence images of anaphase HEK293T cells with CENP-T^ΔC^-dCas9-EGFP targeting after 30min AurKBi treatment, showing examples of EGFP+ mis-segregation

**Supplementary Figure 3.**
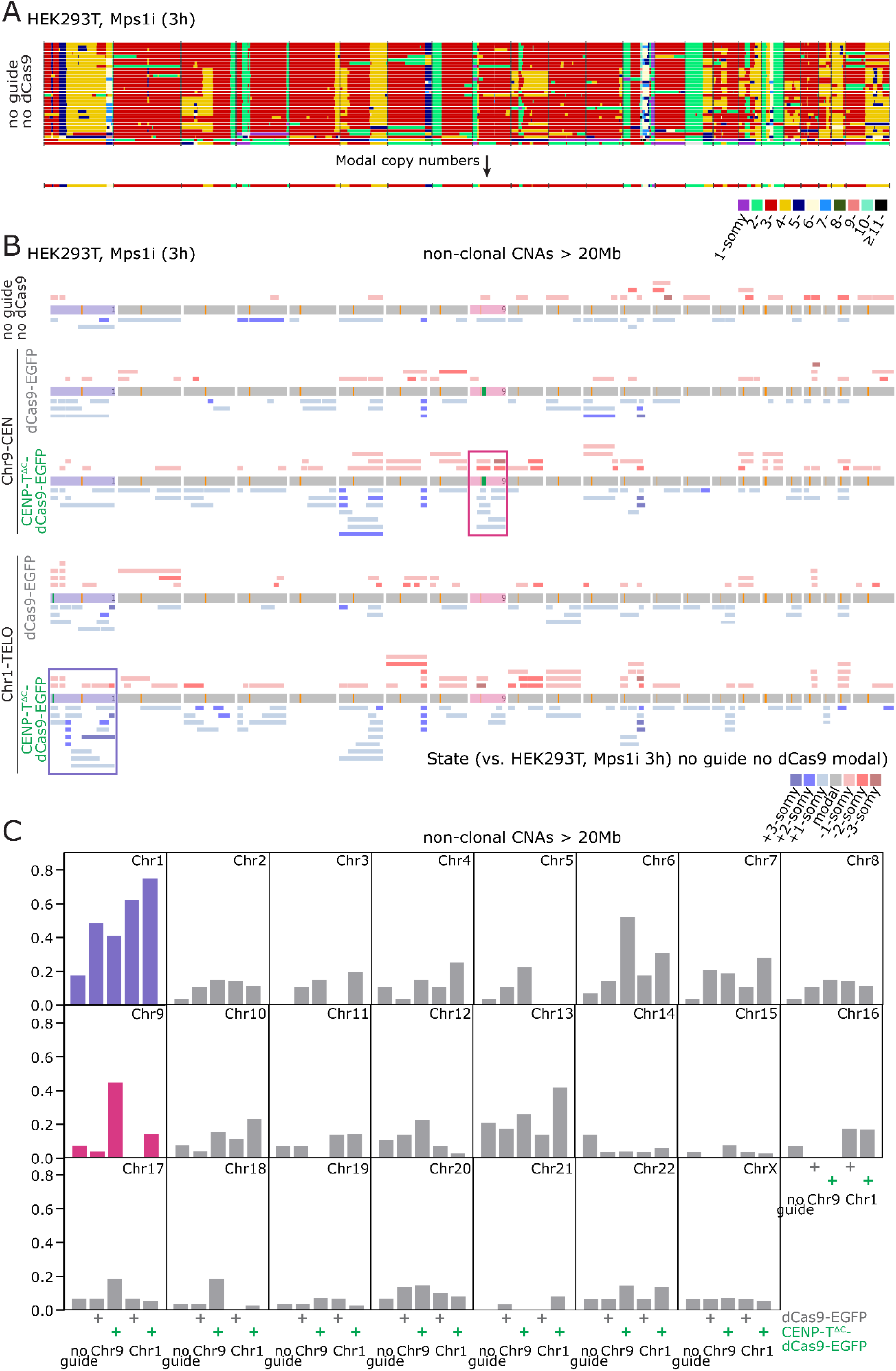
related to Figure 5. **A)** Heatmap showing copy number alterations (CNAs) in single HEK293T cells treated with Mps1i. Heterogeneity is seen at the whole, and partial chromosome level between cells. Median ploidy was calculated across the genome to produce a modal karyotype (lower heatmap). **B)** CNA pileups output from ClonalMasker indicating the CNAs present in each condition not including any clonal or subclonal CNAs present in the Mps1i-only treated cells. Copy number gains are indicated in red (pale red = +1, red = +2, dark red = +3 copies above median reference) and losses in blue (pale blue = −1, blue = −2, dark blue = −3 copies below reference). Orange lines indicate endogenous centromeres and green lines indicate CENP-T^ΔC^ target sites **C)** CNAs >20 MB in size calculated per chromosome per cell for each condition indicated.

**Movie S1: CENP-T^ΔC^-dCas9-nucleated ectopic kinetochores induce prolonged metaphase, and target chromosome mis-segregation**. HEK293T cells targeted with dCas9-CENPT^ΔC^ on chromosome 9 (Chr9-CEN) pausing in metaphase before going through anaphase. DNA is stained in blue (Hoescht) and the targeting foci in green (EGFP). At anaphase onset, the EGFP signals are asymmetrically distributed into the 2 daughter cells. Total length of the movie is 8h.

